# Genetic evidence that lower circulating FSH levels lengthen menstrual cycle, increase age at menopause, and impact reproductive health: a UK Biobank study

**DOI:** 10.1101/028001

**Authors:** Katherine S. Ruth, Robin N. Beaumont, Jessica Tyrrell, Samuel E. Jones, Marcus A. Tuke, Hanieh Yaghootkar, Andrew R. Wood, Rachel M. Freathy, Michael N. Weedon, Timothy M. Frayling, Anna Murray

**Author notes:** Corresponding author: Dr Anna Murray, Genetics of Complex Traits, University of Exeter Medical School, RILD Level 3, Royal Devon & Exeter Hospital, Barrack Road, Exeter, EX2 5DW.

## Abstract

**Study question:** How does a genetic variant altering follicle stimulating hormone (FSH) levels, which we identified as associated with length of menstrual cycle, more widely impact reproductive health?

**Summary answer:** The T allele of the *FSHB* promoter polymorphism (rs10835638) results in longer menstrual cycles and later menopause and, while having detrimental effects on fertility, is protective against endometriosis.

**What is known already:** The *FSHB* promoter polymorphism (rs10835638) affects levels of *FSHB* transcription and, as a result, levels of FSH. FSH is required for normal fertility and genetic variants at the *FSHB* locus are associated with age at menopause and polycystic ovary syndrome (PCOS).

**Study design, size, duration:** We conducted a genetic association study using cross-sectional data from the UK Biobank.

**Participants/materials, setting, methods:** We included white British individuals aged 40–69 years in 2006–2010, included in the May 2015 release of genetic data from UK Biobank. We conducted a genome-wide association study (GWAS) in 9,534 individuals to identify genetic variants associated with length of menstrual cycle. We tested the FSH lowering T allele of the *FSHB* promoter polymorphism (rs10835638) for associations with 29 reproductive phenotypes in up to 63,350 individuals.

**Main results and the role of chance:** In the GWAS for menstrual cycle length, only variants near the *FSHB* gene reached genome-wide significance (*P*<5×10^−8^). The FSH-lowering T allele of the *FSHB* promoter polymorphism (rs10835638G>T; MAF 0.16) was associated with longer menstrual cycles (0.16 s.d. (approx. 1 day) per minor allele; 95% CI 0.12–0.20; *P*=6×10^−16^), later age at menopause (0.13 years per minor allele; 95% CI 0.04-0.22; *P*=5.7×10^−3^), greater female nulliparity (OR=1.06; 95% CI 1.02-1.11; *P*=4.8×10^−3^) and lower risk of endometriosis (OR=0.79; 95% CI 0.69–0.90; *P*=4.1×10^−4^). The FSH-lowering T allele was not associated more generally with other reproductive illnesses or conditions and we did not replicate associations with male infertility or PCOS.

**Limitations, reasons for caution:** The data included might be affected by recall bias. Women with a cycle length recorded were aged over 40 and were approaching menopause, however we did not find evidence that this affected the results. Many of the illnesses had relatively small sample sizes and so we may have been under-powered to detect an effect.

**Wider implications of the findings:** We found a strong novel association between a genetic variant that lowers FSH levels and longer menstrual cycles, at a locus previously robustly associated with age at menopause. The variant was also associated with nulliparity and endometriosis risk. We conclude that lifetime differences in circulating levels of FSH between individuals can influence menstrual cycle length and a range of reproductive outcomes, including menopause timing, infertility, endometriosis and PCOS.

## Introduction

Follicle stimulating hormone (FSH) is a key pituitary expressed hormone, which stimulates maturation of oocytes and is a biomarker of ovarian reserve. FSH is a heterodimer comprised of a hormone specific β-chain (FSHB) associated with an α-chain shared by other members of the glycoprotein hormone family (Nagirnaja, et al., 2010). The anterior pituitary produces FSH with transcription of FSHB rate-limiting for FSH production. FSH stimulates target cells by binding to the FSH receptor (FSHR), a G-protein coupled receptor (Fan and Hendrickson, 2005), promoting follicle maturation and oestrogen production in women, and Sertoli cell proliferation and spermatogenesis in males (Nagirnaja, et al., 2010).

Rare mutations in the *FSHB* gene cause truncation of the FSHB protein and result in hypogonadism and primary amenorrhea in females (Kottler, et al., 2010, Layman, et al., 1997, Matthews and Chatterjee, 1997) and, in a male, delayed puberty with azoospermia (Phillip, et al., 1998). Mouse models suggest that FSH is required for normal fertility. Female *Fshb* knockout mice are infertile and fail to complete normal folliculogenesis while male knockouts remain fertile but have reduced sperm counts, and infertility is observed in both male and female transgenic mice overexpressing human FSH (Kumar, et al., 1999, Kumar, et al., 1997).

A polymorphism in the promotor of *FSHB* (rs10835638G>T) -211 bp upstream of the FSHB transcription start site is associated with reduced FSHB *in vitro* and in human genetic studies. *In vitro*, the T allele of the promotor polymorphism reduces expression of a luciferase reporter gene (Hoogendoorn, et al., 2003) and decreases *FSHB* transcription in gonadotrope cells as a result of reduced LHX3 homeodomain transcription factor binding (Benson, et al., 2013). The T allele of rs10835638 is associated with lower FSH levels in males and females, and is associated with infertility in males (Grigorova, et al., 2008, Grigorova, et al., 2010, La Marca, et al., 2013, Ruth, et al., 2015, Schuring, et al., 2013, Simoni and Casarini, 2014, Tuttelmann, et al., 2012). Genetic association studies have identified signals at the FSHB locus associated with age at menopause (Day, et al., 2015, Stolk, et al., 2012), polycystic ovary syndrome (PCOS) (Hayes, et al., 2015) and levels of luteinising hormone (LH) (Hayes, et al., 2015, Ruth, et al., 2015).

In the first genome-wide association study for menstrual cycle length, we identified the *FSHB* locus as the only signal associated with this trait. Using the unique resource of the UK Biobank (Allen, et al., 2014), we show that a common genetic variant known to alter FSH levels impacts a wide range of traits important to reproductive health, including fertility, endometriosis and menstrual cycle length.

## Methods

### Source of data

The UK Biobank includes 503,325 people aged 40–69 years recruited in 2006–2010 from across the UK (Allen, et al., 2014). We analysed data from the May 2015 interim release of imputed genetic data from UK Biobank which contains 73,355,667 SNPs, short indels and large structural variants in 152,249 individuals (http://www.ukbiobank.ac.uk/wp-content/uploads/2014/04/imputation_documentation_May2015.pdf). UK Biobank invited 9.2 million people to participate, giving a response rate of 5.47% (Allen, et al., 2012). Participants were registered with the NHS and lived within 25 miles of one of the 22 assessment centres. Participants answered detailed questions about themselves, had measurements taken and provided samples.

### Phenotypes

We derived reproductive phenotypes from the UK Biobank data (Supplementary methods). Continuous phenotypes were age at birth of first and last child (females only), age at menarche, age at natural menopause, length of menstrual cycle, number of live births and number of children fathered. To test assumptions of linearity, we analysed the binary outcomes early menarche (lower 5% tail), early menopause (20–44 years), long menstrual cycle (>31 days), short menstrual cycle (<20 days), and multiple pregnancy loss (>1 case).

We defined two infertility-related binary phenotypes; never pregnant (females) and never fathered a child (males). We analysed medical conditions as binary outcomes comparing people reporting a condition (case) with those who did not (control). Medical conditions included dysmenorrhea, endometriosis, fibroids, irregular menstrual cycles, menopausal symptoms, menorrhagia, ovarian cysts, polycystic ovary syndrome (PCOS), uterine polyps, vaginal/uterine prolapse and breast, endometrial and ovarian cancer. As more general indicators of gynaecological health, we included the medical interventions bilateral oophorectomy or hysterectomy in our analysis. Summary statistics for the reproductive traits are presented in Tables 1 and 2.

**Table 1.** Description of cohort of unrelated individuals for continuous outcome measures.

| Phenotype | N | Min | Max | Mean | S.D. | Lower quartile | Median | Upper quartile |
| --- | --- | --- | --- | --- | --- | --- | --- | --- |
| Age at first birth (years)<sup>1</sup> | 43,066 | 10 | 50 | 25.1 | 4.6 | 22 | 25 | 28 |
| Age at last birth (years)<sup>1</sup> | 43,008 | 15 | 50 | 30.0 | 4.8 | 27 | 30 | 33 |
| Age at menarche (years)<sup>1</sup> | 61,306 | 9 | 17 | 12.9 | 1.6 | 12 | 13 | 14 |
| Age at natural menopause (years)<sup>1</sup> | 27,996 | 18 | 65 | 49.9 | 4.5 | 48 | 50 | 53 |
| Length of menstrual cycle (days)<sup>1</sup> | 8,870 | 7 | 300 | 26.8 | 6.2 | 25 | 28 | 28 |
| Number of children fathered<sup>2</sup> | 56,508 | 0 | 28 | 1.8 | 1.2 | 1 | 2 | 2 |
| Number of live births<sup>1</sup> | 63,306 | 0 | 22 | 1.8 | 1.2 | 1 | 2 | 2 |

<sup>1</sup>Females only; <sup>2</sup>Males only.

**Table 2.**
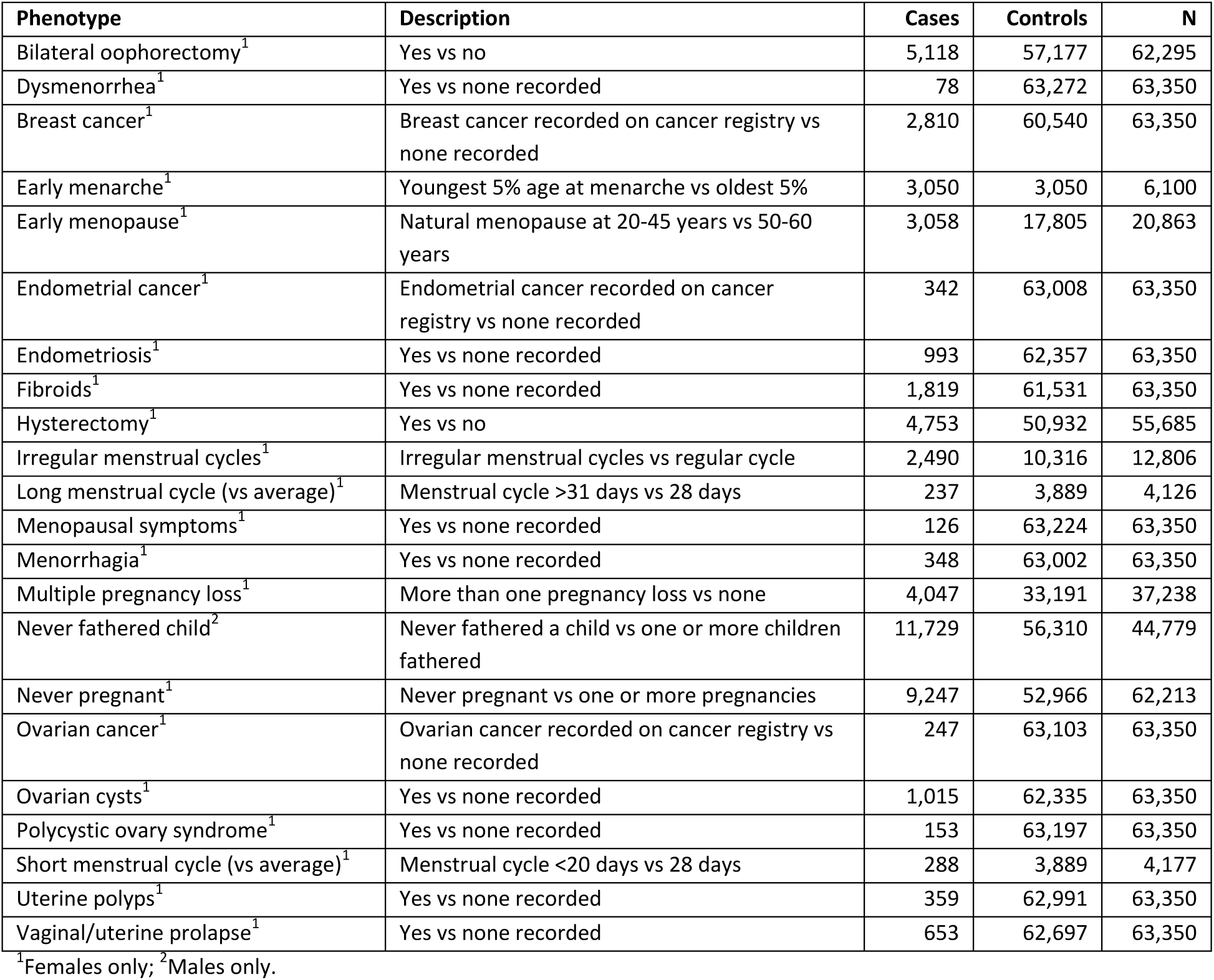
Number of people included in binary outcome measures.

### Participants

In our analysis, we included individuals who both self-identified as white British and were confirmed as ancestrally Caucasian by UK Biobank from genetic information (n=128,266). We calculated principal components (PCs) for inclusion as covariates in our analyses using FlashPCA (Abraham and Inouye, 2014). PCs were calculated in 120,286 unrelated participants (as identified by UK Biobank) based on 95,535 independent, directly genotyped SNPs (pairwise r^2^ <0.1). These SNPs had MAF≥2.5% and missing-ness<1.5% across all participants in the May 2015 interim release of genetic data, and had HWE *P*>1×10^−6^ within the white British participants.

### GWAS of length of menstrual cycle

We conducted a genome-wide association study (GWAS) to identify genetic variants associated with length of menstrual cycle (n=9,534) using BOLT-LMM to account for relatedness and population structure (Loh, et al., 2015). We transformed length of menstrual cycle by adjusting for recruitment centre and age at recruitment prior to inverse-normalisation, and performed association testing while adjusting for genotype chip. We filtered results on imputation quality>0.4, Hardy-Weinberg equilibrium *P*>1×10^−5^, and minor allele frequency >0.1%, resulting in approximately 14 million variants that were tested.

### Testing for associations of the FSHB promoter polymorphism with reproductive phenotypes

We tested the FSH lowering T allele of the *FSHB* promoter polymorphism (rs10835638) for associations with reproductive phenotypes (total n=120,286). SNP rs10835638 was well-imputed in the data (imputation quality 0.995; HWE *P*=0.16; missing rate=0.3%). All analyses were carried out in males or females as appropriate (based on self-defined sex) using Stata (v13).

For continuous phenotypes, we transformed the phenotype by adjusting for recruitment centre, age at recruitment and first five PCs prior to inverse-normalisation. We performed linear regression of imputed minor-allele dosages at SNP rs10835638 on transformed phenotype with genotyping chip as a covariate. We carried out sensitivity analysis of the effect of different transformations, e.g. inverse normalising the trait prior to calculating the residuals, however this did not materially affect our results. For age at menopause and age at menarche, we also ran analysis using the *ReproGen* definition of the phenotype (untransformed age at menopause between 40 and 60 years not adjusted for age, untransformed age at menarche) to allow comparisons with published data (Day, et al., 2015, Perry, et al., 2014, Stolk, et al., 2012). For binary outcomes, we performed logistic regression of the phenotype on minor-allele dosages at SNP rs10835638 including the first five PCs, recruitment centre, age at recruitment and genotyping chip as covariates.

## Results

### A common allele in the FSHB gene, known to lower FSH levels, is associated with longer length of menstrual cycle

The FSH lowering T allele of the FSHB promoter polymorphism (rs10835638G>T; MAF 0.16) was associated with longer menstrual cycles (0.16 s.d. (approx. 1 day) per minor allele; 95% CI 0.12–0.20; *P*=6×10^−16^). Of the reproductive traits tested, length of menstrual cycle was the most strongly associated with rs10835638 (Figure 1). The SNP was also associated with cycle length when we dichotomised into women reporting a cycle length of less than 28 days compared to those reporting the average length of 28 days (OR=0.70; 95% CI 0.54–0.90; *P*=5.1×10^−3^) (Figure 1). There was no evidence for an association with longer than 28 days compared to the average (OR=1.16; 95% CI 0.92–1.47; *P*=0.21). We also conducted analysis excluding cycle length values more than 3 s.d. from the mean (n=8,792; mean 26.6; s.d. 3.33; range 9-45) and results remained consistent.

**Figure 1.**
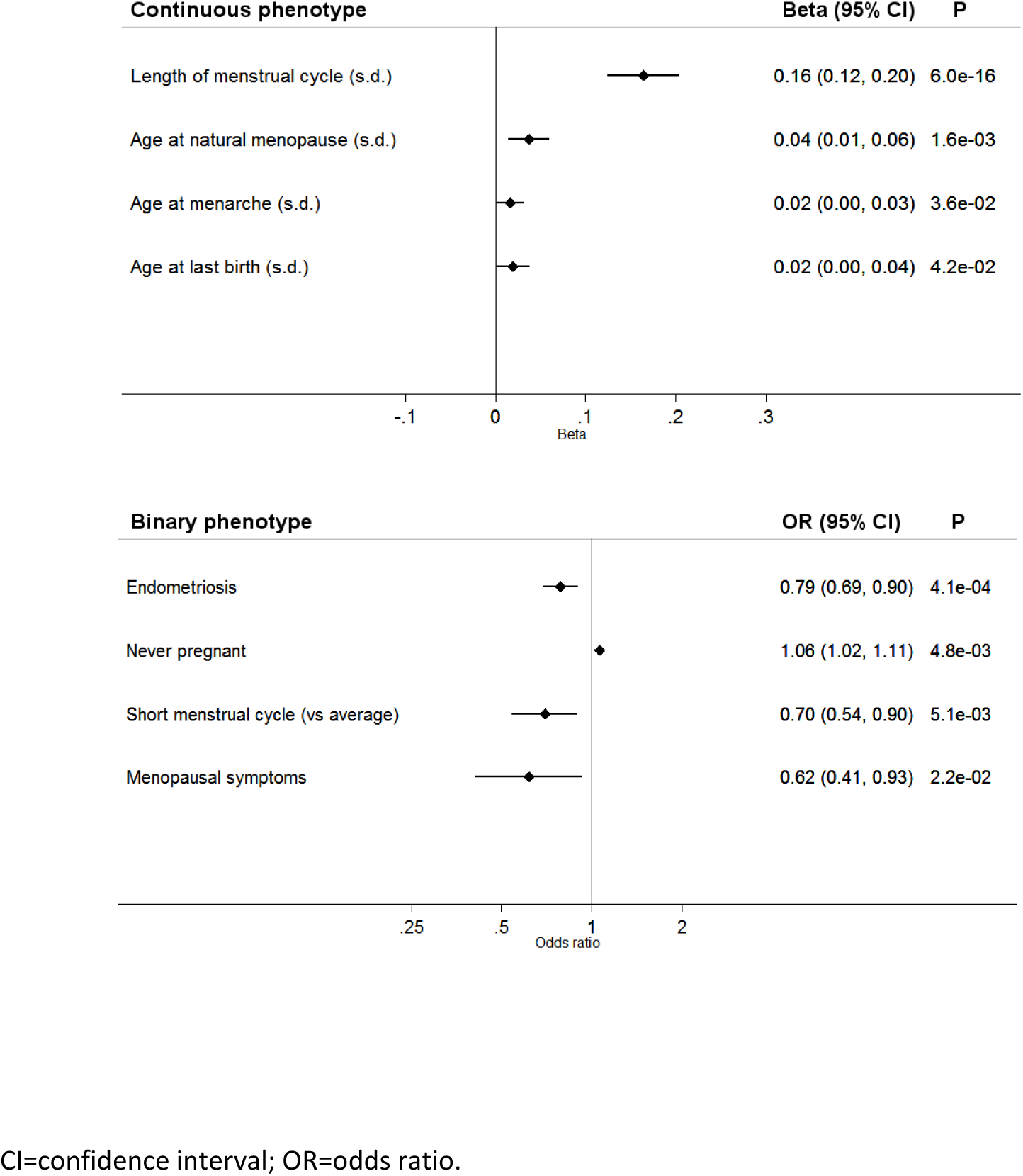
Phenotypes associated (p<0.05) with the FSH lowering allele of rs10835638G>T. For continuous variables, effects (beta) are in standard deviations of the inverse-normally transformed variable to enable effect size comparisons.

Variants in or near the *FSHB* gene were the only ones that reached genome-wide significance in the GWAS for menstrual cycle length (Figure 2). The strongest association was for rs564036233G>GA, a 1 b.p. insertion which was associated with longer cycles by 1 day (0.16 s.d.) per minor allele (95% CI 0.12–0.20; *P*=1.30×10^−16^). The rs564036233 variant is in strong linkage disequilibrium (LD) with the promoter polymorphism rs10835638 (r^2^=0.82) and conditional analysis indicated that rs564036233 and rs10835638 represent the same signal.

**Figure 2.**
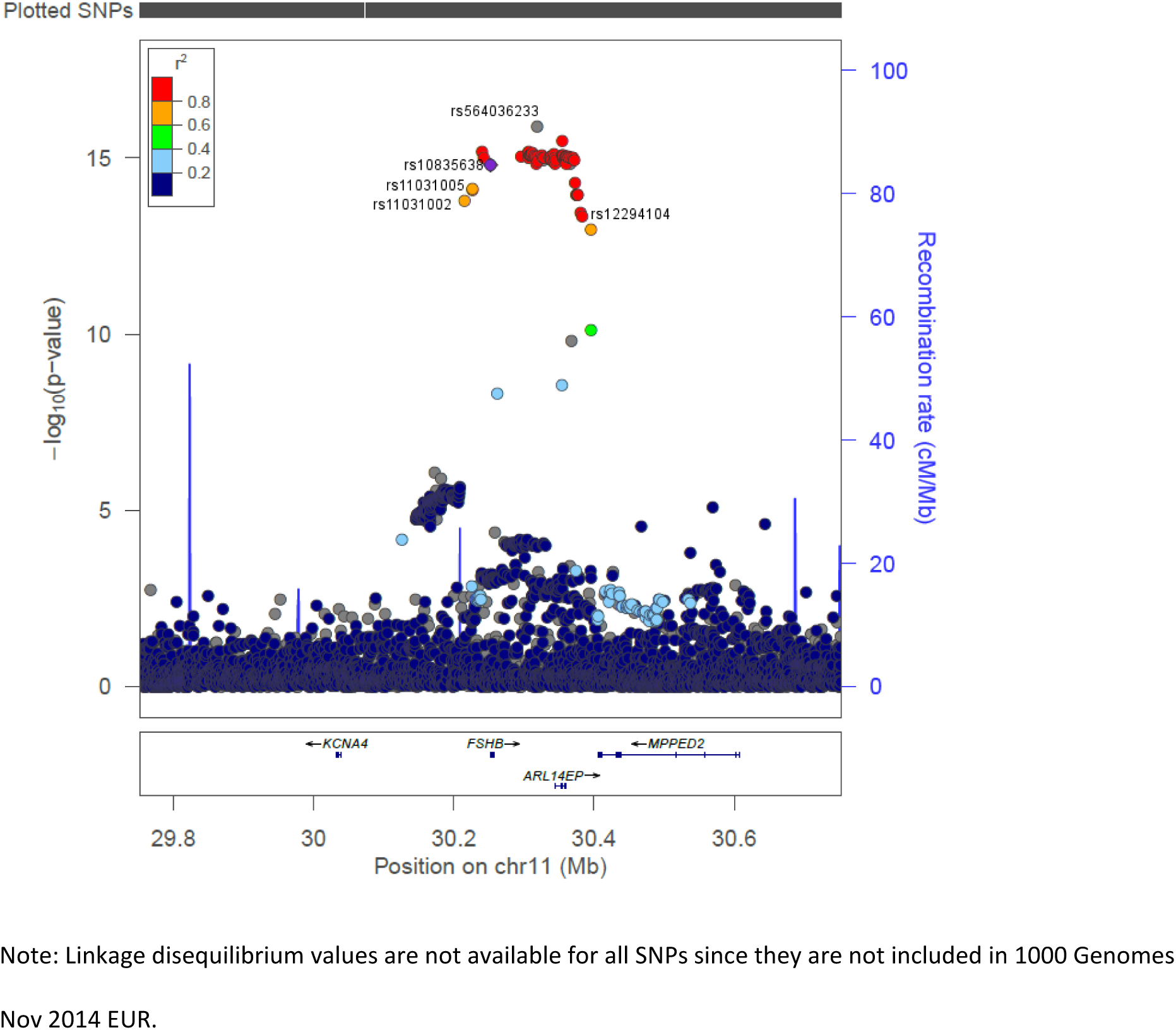
LocusZoom plot showing variants associated with length of menstrual cycle. The most strongly associated variant for cycle length is rs564036233. Linkage disequilibrium (1000 Genomes Nov 2014 EUR) shown is with rs10835638, the FSHB promoter polymorphism. Other SNPs indicated were the variants most significantly associated with FSHB (rs11031005) and LH (rs11031002) in a GWAS of hormone levels (Ruth, et al., 2015), and with age at natural menopause (rs12294104) in a meta-analysis (Stolk, et al., 2012).

### The FSHB allele associated with longer cycle length is associated with later menopause

The FSH lowering T allele of rs10835638G>T was associated with later age at menopause in UK Biobank (0.13 years per minor allele (*ReproGen* definition); 95% CI 0.04-0.22; *P*=5.7×10^−3^). There was no association between rs10835638G>T and menopause age when we dichotomised the phenotype into early menopause compared to later menopause (Table 3). The *FSHB* locus is known to be associated with timing of menopause: in *ReproGen* consortium data, the signal at this locus (rs12294104) increases age at menopause by 0.23 years (95% CI 0.16-0.29; *P*=1.5×10^−11^) (Stolk, et al., 2012). Later menopause is associated with later age at last birth and rs10835638G>T was also associated with later age at last birth (0.02 s.d. (approx. 0.1 years) per T allele; 95% CI 0.00-0.04; *P*=4.2×10^−2^).

**Table 3.** Associations with the FSH lowering T allele of rs10835638G>T. For continuous variables, effects (beta) are in standard deviations of the inverse-normally transformed variable to enable effect size comparisons.

| Phenotype | Statistic | Effect(95% CI) | SE | P |
| --- | --- | --- | --- | --- |
| Length of menstrual cycle (s.d.) | Beta | 0.16(0.12,0.20) | 0.02 | 6.0E-16 |
| Endometriosis | OR | 0.79(0.69,0.90) | 0.05 | 4.1E-04 |
| Age at natural menopause (s.d.) | Beta | 0.04(0.01,0.06) | 0.01 | 1.6E-03 |
| Never pregnant | OR | 1.06(1.02,1.11) | 0.02 | 4.8E-03 |
| Short menstrual cycle (vs average) | OR | 0.70(0.54,0.90) | 0.09 | 5.1E-03 |
| Menopausal symptoms | OR | 0.62(0.41,0.93) | 0.13 | 2.2E-02 |
| Age at menarche (s.d.) | Beta | 0.02(0.00,0.03) | 0.01 | 3.6E-02 |
| Age at last birth (s.d.) | Beta | 0.02(0.00,0.04) | 0.01 | 4.2E-02 |
| Age at first birth (s.d.) | Beta | 0.02(0.00,0.03) | 0.01 | 7.9E-02 |
| Number of live births (s.d.) | Beta | -0.01(-0.03,0.00) | 0.01 | 8.1E-02 |
| Never fathered a child | OR | 1.03(0.99,1.08) | 0.02 | 1.2E-01 |
| Early menopause | OR | 0.95(0.88,1.02) | 0.04 | 1.6E-01 |
| Early menarche | OR | 0.94(0.85,1.04) | 0.05 | 2.1E-01 |
| Fibroids | OR | 0.94(0.86,1.03) | 0.04 | 2.1E-01 |
| Long menstrual cycle (vs average) | OR | 1.16(0.92,1.47) | 0.14 | 2.1E-01 |
| Polycystic ovary syndrome | OR | 1.18(0.88,1.59) | 0.18 | 2.7E-01 |
| Ovarian cysts | OR | 0.94(0.83,1.07) | 0.06 | 3.6E-01 |
| Number of children fathered (s.d.) | Beta | 0.01(-0.01,0.02) | 0.01 | 4.1E-01 |
| Menorrhagia | OR | 0.92(0.74,1.13) | 0.10 | 4.2E-01 |
| Irregular menstrual cycles | OR | 0.97(0.89,1.06) | 0.04 | 4.6E-01 |
| Multiple pregnancy loss | OR | 0.98(0.91,1.04) | 0.03 | 4.6E-01 |
| Dysmenorrhea | OR | 0.87(0.56,1.38) | 0.20 | 5.6E-01 |
| Breast cancer | OR | 1.02(0.95,1.10) | 0.04 | 6.4E-01 |
| Ovarian cancer | OR | 0.94(0.74,1.21) | 0.12 | 6.4E-01 |
| Vaginal/uterine prolapse | OR | 0.97(0.83,1.13) | 0.08 | 6.7E-01 |
| Uterine polyps | OR | 0.98(0.80,1.20) | 0.10 | 8.6E-01 |
| Endometrial cancer | OR | 1.00(0.81,1.23) | 0.11 | 9.7E-01 |

CI=confidence interval; OR=odds ratio; s.d.=standard deviations.

### Longer cycle length is not a general feature of alleles associated with later age at menopause

We next tested the role of all other genetic variants associated with age at menopause. Only one of the other 55 published age at menopause signals was nominally associated with cycle length (p>0.05): rs10734411 was associated at p=0.005 (Day, et al., 2015, Perry, et al., 2014, Stolk, et al., 2012). For the 56 published menopause SNPs there was no correlation between the published effect estimates for age at menopause and the effect estimates from the GWAS for menstrual cycle length (R=0.064, p=0.63) (Figure 3). The *FSHB* SNP was an outlier in this plot, but removing it did not substantially affect the correlation (R=-0.027; p=0.84).

**Figure 3.**
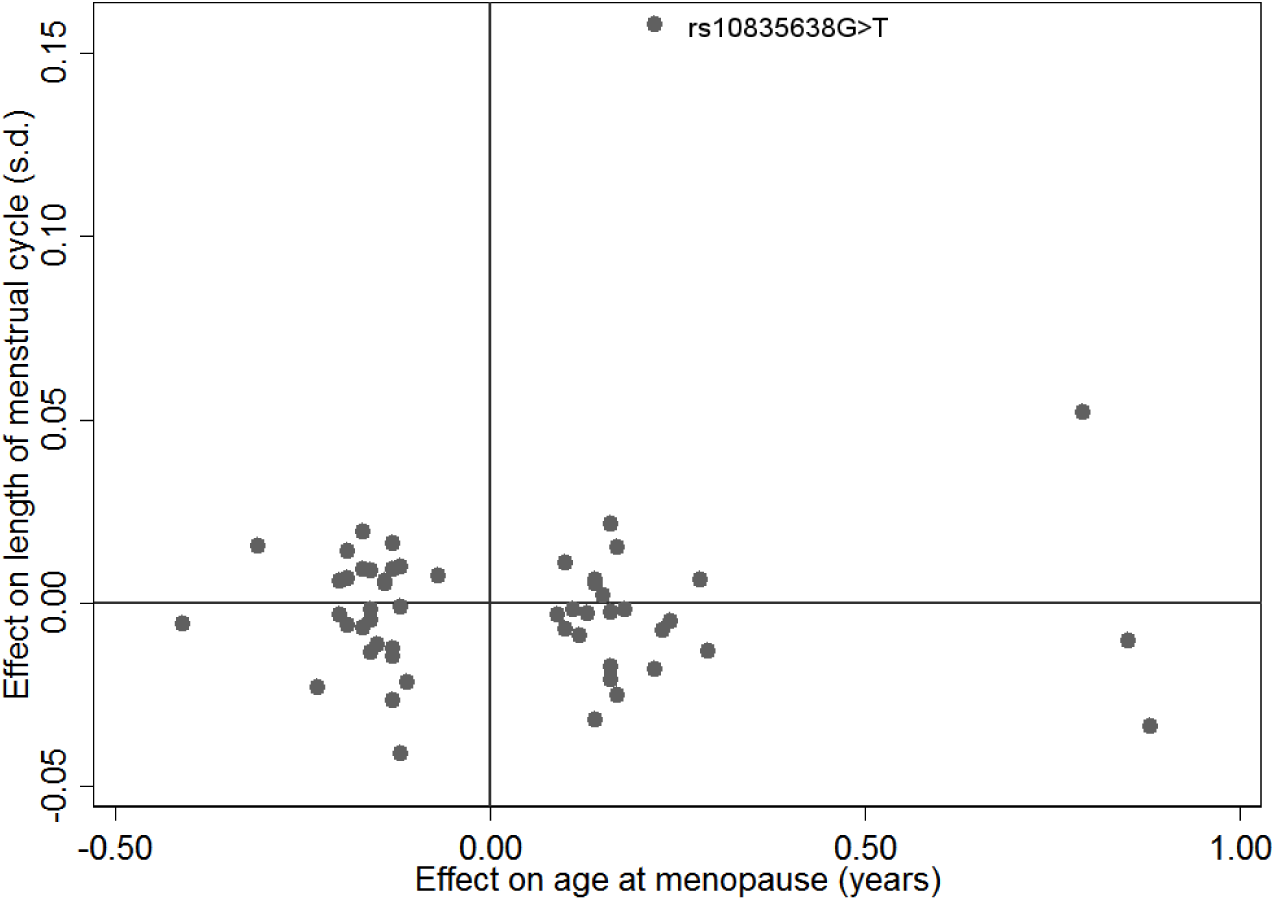
Comparison of the published effect size of the 56 known age at menopause variants (Perry, et al., 2014, Stolk, et al., 2012) and their effect size in the GWAS for menstrual cycle length. There was no significant correlation between the effects on age at menopause and cycle length (R= 0.064, p=0.63). The FSHB promoter polymorphism (rs10835638G>T) is indicated.

### The FSHB allele associated with lower FSH is also associated with an indicator of female infertility

The FSH lowering T allele of the *FSHB* promoter polymorphism (rs10835638G>T) was associated with female nulliparity, i.e. greater odds of never being pregnant (OR=1.06; CI 1.02-1.11*; P*=4.8×10^−3^) (Figure 1). The FSH lowering allele was not associated with other possible indicators of female infertility (later age at first birth and fewer live births) or male infertility (number of children fathered) (p>0.05) (Table 3).

### The FSHB allele associated with higher FSH is also associated with higher odds of endometriosis and surgical intervention

The more common G allele was associated with increased odds of endometriosis (OR=1.27; CI 1.11-1.45; *P*=4.1×10^−4^) (Figure 1). Of the seven published GWAS variants associated with endometriosis risk (Nyholt, et al., 2012), the variant on chromosome 12 was nominally associated with cycle length, with the allele associated with an increased risk of endometriosis also associated with shorter cycles (p=0.02).

The G allele of rs10835638G>T was also associated with increased odds of having the medical interventions bilateral oophorectomy (OR=1.12; 95% CI 1.06-1.19; *P*=1.4×10^−4^) and hysterectomy (OR=1.13; 95% CI 1.06-1.20; *P*=1.0×10^−4^), which are used as treatments for a range of gynaecological conditions including endometriosis.

### The common FSHB variant, associated with FSH levels, is not associated with reproductive traits more generally

There was no consistent evidence that the *FSHB* variant was associated with age at menarche. There was a 0.03 year increase in age at menarche (*ReproGen* definition) per T allele of rs10835638G>T (95% CI 0.01-0.05; *P*=1.4×10^−2^) and the binary phenotype of early menarche was associated at p>0.05 (Table 3). None of 122 published GWAS signals for menarche (Perry, et al., 2014) were associated with length of menstrual cycle at p<0.008.

The *FSHB* promoter polymorphism (rs10835638G>T) was not associated with other reproductive illnesses or conditions at *P*<0.05 (Table 3), except for menopausal symptoms (OR=0.62; 95% CI 0.41-0.93; p=0.02) (Figure 1). No associations were found with dysmenorrhea, fibroids, irregular menstrual cycles, menorrhagia, multiple pregnancy loss, ovarian cysts, PCOS, uterine polyps or vaginal/uterine prolapse, or with female breast, ovarian or endometrial cancer.

## Discussion

In the first GWAS of menstrual cycle length we found a strong association between an FSH lowering, likely functional, variant in the *FSHB* promoter and longer cycles (Benson, et al., 2013, Grigorova, et al., 2008, Grigorova, et al., 2010, Hoogendoorn, et al., 2003, La Marca, et al., 2013, Ruth, et al., 2015, Simoni and Casarini, 2014, Tuttelmann, et al., 2012). This locus has been previously robustly associated with age at menopause in the *ReproGen* GWAS of menopause timing (Day, et al., 2015, Stolk, et al., 2012) and the allele associated with longer cycle length is associated with later age at menopause. We did not observe associations for the majority of age at menopause GWAS signals with length of menstrual cycle, including the four signals with effects of over one-third of a year per allele on menopause timing, implying that the association is specific to *FSHB:* Either FSHB has independent effects on both cycle length and menopause, or changes in cycle length are causally influencing menopause timing.

Our results are consistent with the observed epidemiological relationship between longer menstrual cycles and later age at menopause (Kaczmarek, 2007, Whelan, et al., 1990). It is possible that there is a biological limit on the lifetime number of menstrual cycles, hence women with longer cycles would have later menopause. Alternatively, they may have reduced follicle recruitment per cycle, depleting their ovarian reserve more slowly. Women with longer cycles have more waves of folliculogenesis during each cycle (Baerwald, et al., 2003, Baerwald, et al., 2012) but may recruit fewer antral follicles per wave. Oocyte loss due to ovulation is unlikely to be driving the relationship, since this contributes much less to overall oocyte depletion than atresia, there is no robust evidence that preventing ovulation by use of the combined oral contraceptive pill influences menopause timing (Ayatollahi, et al., 2003, de Vries, et al., 2001, Dorjgochoo, et al., 2008, Gold, et al., 2001, Gold, et al., 2013, Kaczmarek, 2007, OlaOlorun and Lawoyin, 2009, Palmer, et al., 2003, Pokoradi, et al., 2011, Stepaniak, et al., 2013, van Noord, et al., 1997) and both longer and shorter cycles are more likely to be anovulatory (Mihm, et al., 2011). More work is needed to understand the molecular mechanism that explains the association between cycle length and menopause timing.

The FSH-reducing allele was associated with nulliparity, indicating increased female infertility. Although we were unable to distinguish those unable to have children from those not wishing to, nulliparious women will be enriched for both female and male factor infertility. The FSH-lowering allele has previously been found to be associated with male infertility (Grigorova, et al., 2008, Grigorova, et al., 2010, Simoni and Casarini, 2014, Tuttelmann, et al., 2012), but we found no association with males who had never fathered a child suggesting a female-specific effect. Using nulliparity as a proxy for infertility is unlikely to generate a false-positive association, but may have reduced our power to detect a true association. The relationship between FSH and fertility over a woman’s lifetime may differ from the age-related changes in FSH around menopause. In contrast to our genetic association between lower FSH and infertility, women nearing menopause have higher FSH concentrations, poorer ovarian reserve and decreased fertility (Mihm, et al., 2011, Waller, et al., 1998). FSH is required for follicle development and it is proposed that an FSH threshold is required to achieve ovulation (Kumar, et al., 1999, Kumar, et al., 1997). Ovulation increases with increasing FSH in transgenic mice with FSH levels that increase with age independently of follicle depletion (McTavish, et al., 2007). A high baseline level of FSH, determined by genetic variation, may promote ovulation and explain our association with parity.

The FSH-increasing allele increased the risk of endometriosis in our study. Several GWAS of endometriosis have been performed, however none have reported a signal at the 11p14.1 locus and there was no evidence that the genome-wide significant endometriosis variants were associated with cycle length in our study (Adachi, et al., 2010, Albertsen, et al., 2013, Nyholt, et al., 2012, Painter, et al., 2011, Uno, et al., 2010). Drug treatments for endometriosis aim to prevent ovulation and menstruation, and to stabilise hormone levels, since oestrogens fuel ectopic endometrial growth (Vercellini, et al., 2014). The FSH-increasing allele may similarly stimulate abnormal growth of endometrium. Endometriosis is associated with earlier menopause (Pokoradi, et al., 2011, Yasui, et al., 2011) and shorter menstrual cycles (Vercellini, et al., 2014), consistent with our findings. The FSH-increasing variant associated with increased risk of endometriosis was also associated with parity, however endometriosis can cause infertility as a result of endometriotic lesions and chronic pelvic inflammation. Therefore, the association of the *FSHB* polymorphism with infertility appears to be independent of the association with endometriosis.

We found a modest association of the FSH-lowering allele with increased age at menarche, but the published age at menarche GWAS signals were not associated with length of menstrual cycle. The closest GWAS menarche signal to *FSHB* (rs16918636) is 1.13Mb away and is not in LD (r^2^=0.001) with the *FSHB* promoter polymorphism SNP (Perry, et al., 2014). Although FSH is important for normal puberty, the role of variation in baseline FSH levels on puberty timing is uncertain.

UK Biobank recruited individuals over 40 years old, and many of the women still cycling will be approaching menopause, however if the association with cycle length was being driven by peri-menopausal changes we would expect all menopause-associated variants to be associated with cycle length. We were unable to replicate an association between the FSH-lowering allele and increased odds of PCOS (Hayes, et al., 2015). However, we had only a small number of cases (n=153) limiting our power to detect this association. Other illnesses had relatively small sample sizes and we may have been similarly under-powered. We might have also under-ascertained cases, as most illnesses will be subject to recall bias as they are self-reported and collected retrospectively, while controls might include people not reporting an illness.

Our study provides evidence that a likely functional variant in the *FSHB* promoter is strongly associated with longer menstrual cycles, and to a lesser extent with female infertility and lower risk of endometriosis. There is considerable evidence that the T allele of the *FSHB* promoter polymorphism decreases FSH levels (Benson, et al., 2013, Grigorova, et al., 2008, Grigorova, et al., 2010, Hoogendoorn, et al., 2003, La Marca, et al., 2013, Ruth, et al., 2015, Simoni and Casarini, 2014, Tuttelmann, et al., 2012), but it has also been associated with increased LH levels (Hayes, et al., 2015, Ruth, et al., 2015). While we cannot rule out that the variant may be having direct or indirect effects on other hormone levels, a change in FSH is the most likely primary mechanism. In conclusion, we suggest that lower FSH levels result in longer menstrual cycles and as a consequence later menopause and, while having detrimental effects on fertility, are protective against endometriosis.

## Authors’ Roles

A.M. and K.S.R. designed the study, carried out analysis and drafted the article. All authors were involved in designing and performing analysis of the UK Biobank data, revising and approving the manuscript.

## Acknowledgements

We thank Dr A.M. Godwin (DRCOG, MRCGP) for identifying medications influencing length of menstrual cycle.This research has been conducted using the UK Biobank Resource.

